# Capturing the dynamics of genome replication on individual ultra-long nanopore sequence reads

**DOI:** 10.1101/442814

**Authors:** Carolin A Müller, Michael A Boemo, Paolo Spingardi, Benedikt M Kessler, Skirmantas Kriaucionis, Jared T Simpson, Conrad A Nieduszynski

## Abstract

The replication of eukaryotic genomes is highly stochastic, making it difficult to determine the replication dynamics of individual molecules with existing methods. We now report a sequencing method for the measurement of replication fork movement on single molecules by <u>D</u>etecting <u>N</u>ucleotide <u>A</u>nalogue <u>s</u>ignal <u>c</u>urrents on <u>e</u>xtremely long <u>n</u>anopore <u>t</u>races (D-NAscent). Using this method, we detect BrdU incorporated by *Saccharomyces cerevisiae* to reveal, at a genomic scale and on single molecules, the DNA sequences replicated during a pulse labelling period. Under conditions of limiting BrdU concentration, D-NAscent detects the differences in BrdU incorporation frequency across individual molecules to reveal the location of active replication origins, fork direction, termination sites, and fork pausing/stalling events. We used sequencing reads of 20-160 kb, to generate the first whole genome single-molecule map of DNA replication dynamics and discover a new class of low frequency stochastic origins in budding yeast.

## Introduction

Genomic methods have provided insights into DNA replication and genome stability^1-3^. Within a population of cells, these methods mask heterogeneity in both replication origin usage and replication fork dynamics; what happens in each individual cell is difficult to ascertain^4^. A high-throughput single-molecule approach is needed to reveal the heterogeneity in DNA replication dynamics. In addition, such an approach has the potential to identify origins used in organisms with very high levels of heterogeneity, in particular mammalian cells, for which population analysis is less useful.

Current single-molecule techniques to study DNA replication have provided valuable insight, but have limitations. DNA combing relies on antibody detection of nucleotide analogues incorporated on the nascent strand and can be used to determine the pattern of origin activation and fork progression in single molecules^5^. However, this approach is low-throughput and provides limited temporal and spatial resolution: combed molecules are anonymous unless genomic positions are identified by probe hybridization, which is particularly challenging for large metazoan genomes. Alternative methods use of nanochannels to stretch DNA molecules has led to increases in throughput and can help to reveal genomic location, but the temporal and spatial resolution are limited by analogue pulse length and image-based detection, respectively^6, 7^. Recently, *in vitro* systems have been established that use single-molecule imaging to monitor replication protein kinetics on DNA^8,9^. Visualizing individual, fluorescently-tagged proteins provided novel mechanistic insights into replication origin licensing and initiation. However, *in vitro* systems are presently limited to small DNA molecules (replicated from a single origin) and so cannot recapitulate *in vivo* replication dynamics.

Here, we present a nanopore-based sequencing method that can measure replication fork movement by <u>d</u>etecting <u>N</u>ucleotide <u>A</u>nalogue <u>S</u>ignal <u>C</u>urrents on <u>E</u>xtremely long <u>N</u>anopore <u>T</u>races (D-NAscent) in nascent DNA. We demonstrated that currently available nanopore sequencing platforms can reliably distinguish base analogues from natural bases. We have developed software that detects BrdU on individual nanopore sequencing reads: When BrdU is incorporated by replication forks, D-NAscent can detect these regions of incorporation. We demonstrated the power of D-NAscent in *S. cerevisiae* (the eukaryote in which genome replication is best characterized). A pulse-chase experiment revealed the regions replicated during the pulse, providing information comparable to that from DNA combing, but at a genomic scale and with sequence-level information. We validated D-NAscent by comparison to mass-spectrometry and population-level sequencing data. In experiments where BrdU was limiting, we showed that D-NAscent can detect the changes in BrdU incorporation frequency to reveal the direction of replication forks and identify the location of replication origins on individual molecules. Using this approach, we have created a whole-genome profile that reveals the replication fork dynamics and origin firing on each of over 100,000 molecules 20-160 kb in length.

## Results

### Nucleotide analogues produce a distinct signal in nanopore sequencing

The Oxford Nanopore Technologies (ONT) MinION instrument determines a base sequence from the electrical readout produced as a single strand of DNA passes through a protein pore. The MinION samples an ionic current signal over time, where each short DNA sequence within the nanopore can be identified by the magnitude of the signal it produces. To simplify analysis, it is typically assumed that the observed signal only depends on a short fixed-length sequence, which is termed a *k*-mer. The current signal for each *k*-mer can be modelled by a Gaussian distribution. For consistency with the data released by ONT, we used a *k*-mer length of six. We and others have previously demonstrated that signal-level events can distinguish methylated from unmodified bases^10, 11^. To determine whether this platform can also distinguish analogues from natural bases we sequenced DNA substrates where thymidine (at various fixed positions) had been substituted by different synthetic analogues. We observed clear differences in the event distributions between thymidine and 5-bromodeoxyuridine (BrdU) using an earlier sequencing protocol (R9 pore at a sequencing rate of 250bp/s) and the current protocol (R9.5 pore at 450bp/s; **Fig. 1a** and **1b**). (Similar observations were made with the previously available generation of the pore, R7.3 – data not shown.) The shift in signal depended on the particular analogue, the sequence context, and the position of the analogue within the 6-mer; the greatest shift was observed when the analogue was substituted for thymidine at the fourth base from the 5’ end of the 6-mer (**Fig. 1b-e** and **Supplemental Fig. S1**). These observations indicate that MinION sequencing has the potential to detect nucleotide analogues in genomic DNA.

**Figure 1:**
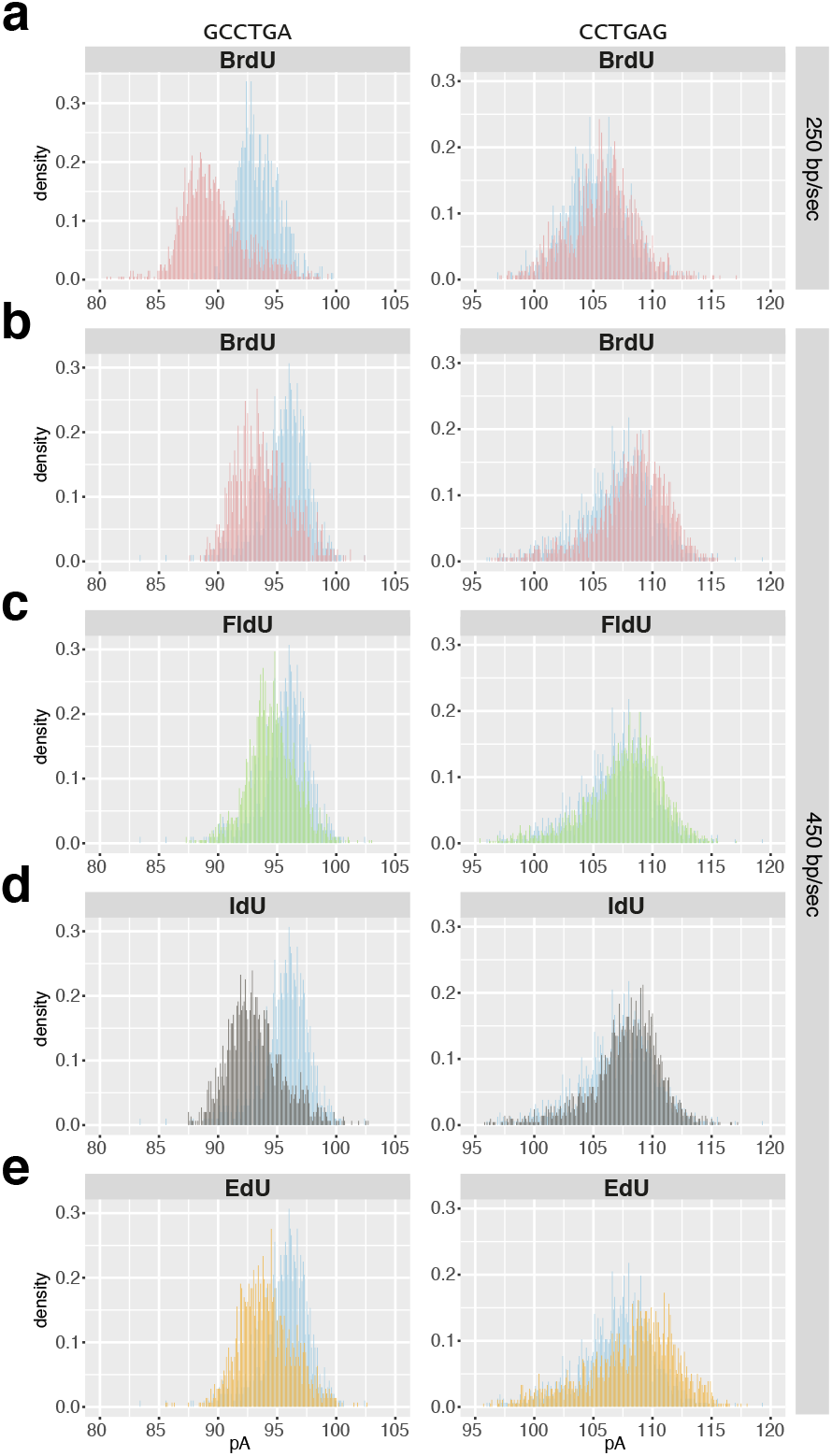
Nanopore sequencing can distinguish thymidine from analogues. For two example 6-mers, each panel shows the distribution of signal events for thymidine (blue) and various analogues: BrdU (**a** and **b**); FldU (**c**); IdU (**d**); and EdU (**e**). The data were generated using the ONT MinION R9 and R9.5 pore with sequencing speeds of 250 bp/s (**a**) or 450 bp/s (**b** – **e**) respectively.

### Identifying the characteristic nanopore signal of BrdU in genomic DNA

We sought to determine the distribution of nanopore signal events for any BrdU-containing 6-mer in genomic DNA. A *Saccharomyces cerevisiae* strain which is dependent upon exogenous thymidine^12^ was grown in various proportions of thymidine and BrdU. Genomic DNA was prepared and analysed by MinION sequencing (see Online Methods). As a control, BrdU incorporation was quantified by mass spectrometry and immunoprecipitation followed by Illumina sequencing (BrdU-seq; **Supplemental Fig. S2**, **S3** and summarised in **Supplemental Table S1**)^13^. The mass spectrometry data revealed that in our five genomic DNA samples the percentage of thymidines substituted by BrdU was 0%, 15%, 26%, 49% and 79%.

Nanopore sequencing signal events were aligned to the genomic reference using nanopolish^14^. These data revealed many thymidine-containing 6-mers where the distribution of signal events was bimodal; while one population matched the ONT model, there was a distinct second population (**Fig. 2a**). The relative proportions of the two populations reflected the concentration of incorporated BrdU. By contrast, 6-mers that did not contain thymidine were mono-modal and matched the ONT model (data not shown). We fit a bimodal Gaussian mixture model to the signal events from the 49% BrdU sample that aligned to each thymidine-containing 6-mer (**Fig. 2b**). We used the Kullback-Leibler (KL) divergence, which measures the average log-likelihood between two probability distributions, to quantify the difference between the ONT model and each of the two fit distributions. One distribution (fit 1) was close to the ONT model (only ∼1% of 6-mers had a KL-divergence >0.5) while the second distribution (fit 2) was farther away from the ONT model (∼62% 6-mers had a KL-divergence >0.5) and corresponded to the BrdU concentration-dependent population (**Fig. 2b** and **c**). We conclude that the second distribution represents the BrdU signal.

**Figure 2:**
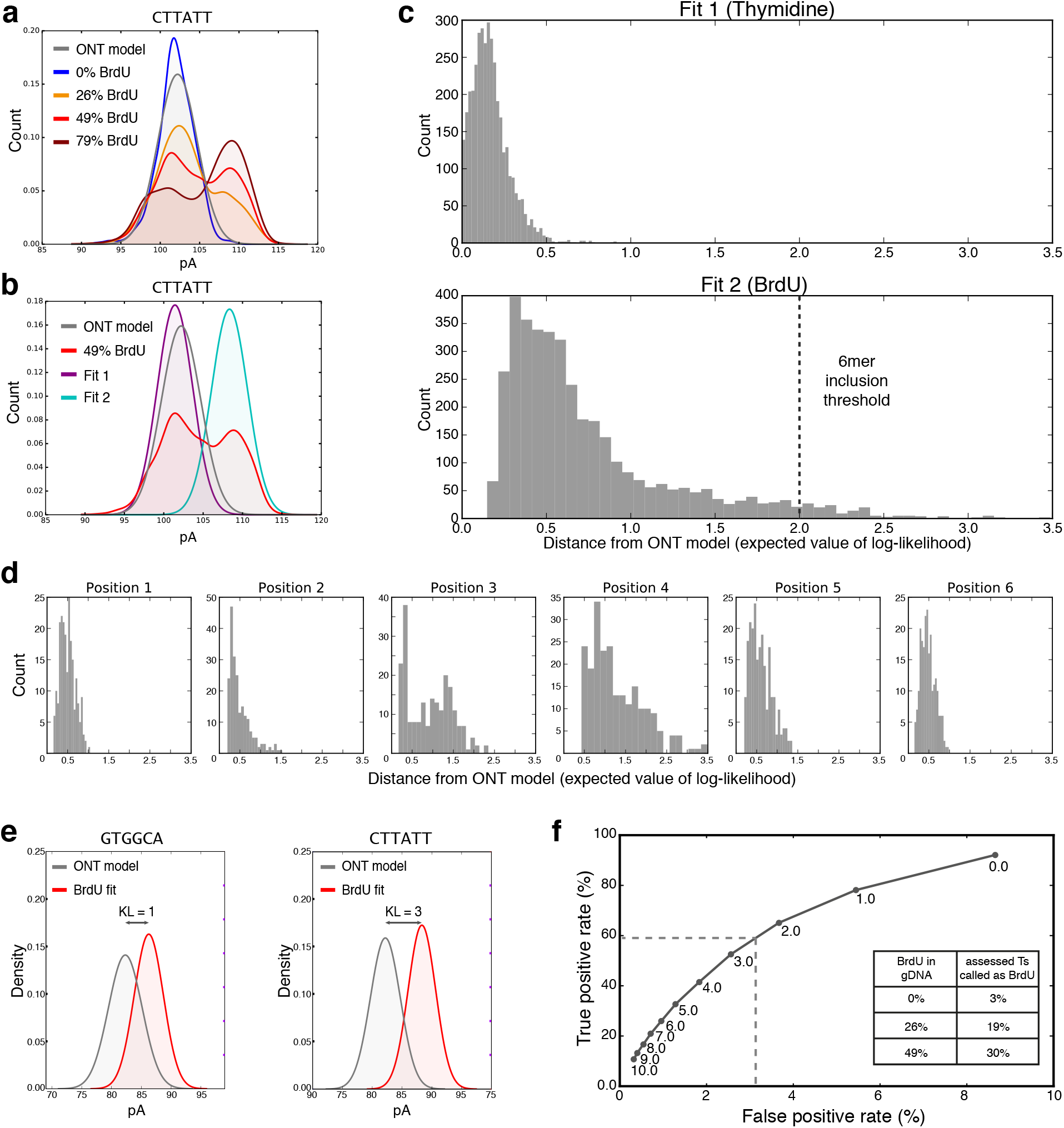
BrdU can be distinguished from thymidine in genomic DNA. (**a**) Signal event distributions for an example 6-mer from yeast genomic DNA containing various concentrations of BrdU (0% - blue; 26% - orange; 49% - red; 79% - crimson) compared to the ONT model (grey). (**b**) Bimodal Gaussian mixture model fit (purple and turquoise) for an example 6-mer from genomic DNA containing 49% BrdU (red). The ONT model is shown in grey. (**c**) Distribution of the KL-divergence between the ONT model and Gaussian fit 1 (upper) or fit 2 (lower) for all thymidine-containing 6-mers. (For detection (Fig. 2e), we make a BrdU call for all 6-mers that have a KL-divergence >2.0; dashed line, lower plot.) (**d**) Distributions as in the lower plot from (c) but for the subset of 6-mers containing just one thymidine; plotted by the position of the thymidine. (**e**) Signal event distributions from the ONT model (thymidine; grey) and from the bimodal Gaussian mixture model fit for BrdU (red). The KL-divergence of the two 6-mers is indicated. (**f**) Receiver operating characteristic (ROC) curve, using all 6-mers that have a KL-divergence >2.0, specifying the true positive and false positive rates for various log-likelihood thresholds of BrdU compared to thymidine (see Online Methods). Numbers near points specify the log-likelihood threshold above which a position in a read is classified as BrdU. The dashed lines demarcate the true and the false positive rates at a log-likelihood threshold >2.5.

We note that even at high BrdU substitution levels 6-mers featuring multiple thymidines gave only a bimodal distribution of signal events (**Fig. 2a**). This is consistent with our previous observation that BrdU predominantly shifts the signal event when present at the fourth base from the 5’ end of the 6-mer (**Fig. 1b** and **Supplemental Fig. S1**). To assess this further, we examined the subset of 6-mers containing a single thymidine and observed the greatest shift in signal event when BrdU is the third or fourth base from the 5’ end (**Fig. 2d**). These data indicate that it will be possible to distinguish BrdU from thymidine in genomic DNA.

### Detection of BrdU incorporated *in vivo*

Detection relied on two thresholds: those 6-mers for inclusion in the model and a certainty above which a position is called as BrdU. Including only those 6-mers where BrdU caused a KL-divergence from the ONT model >2 (*N*=159; **Fig. 2c**) allowed assessment of BrdU incorporation on average every 21 nucleotides across the yeast genome (**Supplemental Fig. S4a**). Each time one of these 6-mers occurred in our sequencing reads, we used a Hidden Markov model (HMM) to compute the log-likelihood that the 6-mer contained a BrdU (**Supplemental Fig. S5**). A position in a read was classified as BrdU if the log-likelihood exceeded a threshold (**Fig. 2e**); this threshold was determined by testing the HMM on unused training data and on an equivalent thymidine-only sample to determine true and false positive rates (**Fig. 2f** and **Supplemental Fig. S4b**). This demonstrated that we can select a threshold that gives a low false positive rate and a reasonably high true positive rate; we set this threshold at log-likelihood >2.5 which gave a true positive rate of ∼60% for a false positive rate of ∼3%. We achieved a similar true positive rate using a DNA sample with an intermediary BrdU concentration (26% incorporation) that was unrelated to the training material.

To further test our detection strategy, we generated hemi-BrdU substituted yeast genomic DNA, by synchronizing a strain prototrophic for thymidine^15^ and passing it through one S phase in media containing BrdU. Material was validated by mass spectrometry (**Supplemental Fig. S2**) and BrdU-seq to reveal any incorporation bias (**Supplemental Fig. S6**). The cell cycle synchrony was confirmed by flow cytometry of DNA content (**Supplemental Fig. S7**). MinION sequencing was performed and positions of BrdU incorporation were determined as described above. As anticipated, we observed reads with either low or predominantly high density of BrdU calls over the entire read, consistent with parental and nascent strands, respectively (**Fig. 3a**). To quantify the frequency of BrdU calls in each read, we fit the number of positive BrdU calls in non-overlapping 2kb windows to a binomial distribution (see Online Methods). This allowed us to compare the number of positive BrdU calls in each window to what we would expect from a typical BrdU-positive window (determined from mass spectrometry data and the true-positive rate). Computing the z-score of positive BrdU calls in each window against this binomial distribution revealed a bimodal density: One population was centered around the mean, indicating these windows had a BrdU frequency consistent with our expectation for a BrdU-positive window; the other population was centered ∼5 s.d. below the mean, indicating these windows had significantly fewer positive BrdU calls than we expect from a BrdU-positive window (**Fig. 3b**). We conclude that these two populations correspond to BrdU-containing windows and thymidine-only windows, respectively. These results indicate that our model can distinguish parental DNA from nascent DNA containing BrdU. We call our method <u>D</u>etecting <u>N</u>ucleotide <u>A</u>nalogue <u>s</u>ignal <u>c</u>urrents on <u>e</u>xtremely long <u>n</u>anopore <u>t</u>races (D-NAscent).

**Figure 3:**
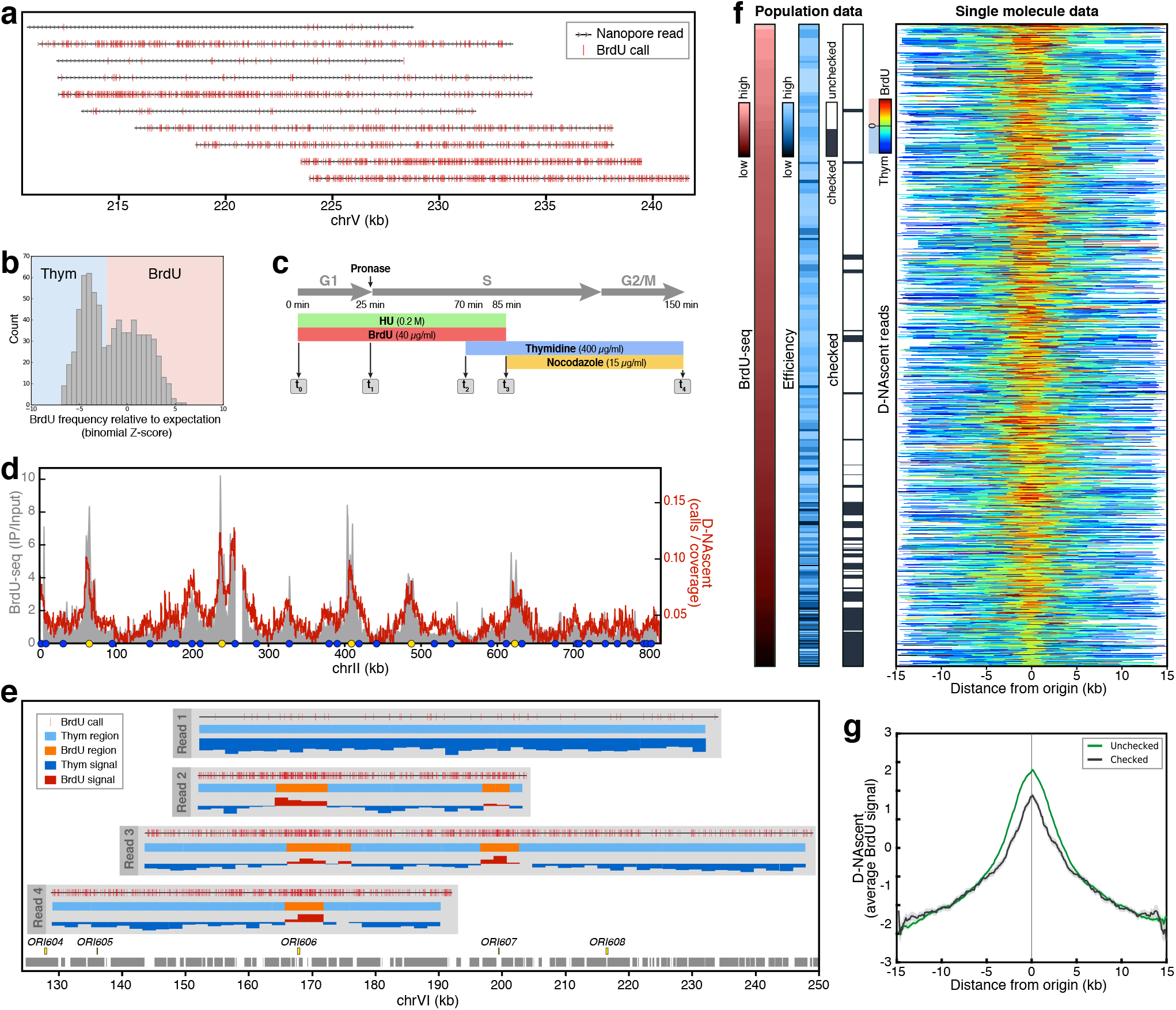
Single-molecule detection of BrdU on nascent DNA. (**a**) Representative nanopore reads (>15 kb) showing BrdU calls in hemi-substituted yeast genomic DNA. Red ticks indicate positive BrdU calls and arrows give the read direction relative to the sacCer3 reference genome. (**b**) The distribution of positive BrdU call frequency measured as a z-score of a binomial distribution for non-overlapping 2 kb windows. (For later analysis we set a binomial z-score threshold >-2 for assigning a window as BrdU positive.) (**c**) Schematic of the experimental strategy for detection of replication origin activity in HU. At each timepoint, samples were taken for mass spectrometry, DNA copy number measurement, BrdU-seq and D-NAscent. (**d**) Comparison of BrdU-seq and an ensemble of D-NAscent data across chromosome II (from timepoint 4). (**e**) Four example nanopore sequencing reads that illustrate BrdU detection on parental and nascent strands mapping to the right end of chromosome VI. Each read shows BrdU calls at individual 6mers (upper track), BrdU-positive 2 kb windows shown (orange; middle track), and the z-score for each window where red bars are above the detection threshold (z-score = −2) and are BrdU-positive (lower track). Confirmed replication origins from OriDB (yellow boxes) and genes (grey boxes) are shown. (**f**) Visualisation of D-NAscent data for 1,325 individual nanopore reads (rows) that span confirmed replication origins (OriDB), ordered by BrdU-seq data. Additional colour bars show population-level data for BrdU-seq, origin activation efficiency^4^ and whether the origin is ‘checked’ by the intra-S phase checkpoint^17^. (**g**) Ensemble BrdU signal from D-NAscent for all ‘unchecked’ (green) and ‘checked’ (black) origins (BrdU z-scores averaged for each column in (**f**); shaded areas show the standard error of the mean).

### Single-molecule detection of replication origin activity in hydroxyurea

DNA synthesis can be slowed by the addition of hydroxyurea (HU), which inhibits ribonucleotide reductase, thereby restricting BrdU incorporation to locations proximal to early activating replication origins^16-18^. To investigate the pattern of origin usage on single-molecules, we released cells synchronously from G1 into S phase in the presence of HU and BrdU. After 60 minutes, cells were chased out of HU, with excess thymidine, into nocodazole (to prevent entry into a second cell cycle; **Fig. 3c**). After completion of S phase, samples were collected for D-NAscent, mass spectrometry, and BrdU-seq. Cell cycle synchrony was assessed by flow cytometry of DNA content (**Supplemental Fig. S7**). BrdU detected in the MinION data was summed across all reads in non-overlapping 100 bp windows to allow comparison to the BrdU-seq data (**Fig. 3d**). This confirmed that an ensemble of our single-molecule data is in good agreement (Pearson correlation coefficient, *R*=0.76) with established short-read methods. Visualising the individual positive BrdU calls on single-molecules suggested that each sequencing read fell into one of two categories: there was either an infrequent number of positive BrdU calls throughout the whole read, or the read contained short patches of frequent positive BrdU calls. (**Fig. 3e**). We conclude that these reads likely correspond to parental and nascent strands, respectively. For each individual read, we assessed non-overlapping 2 kb windows and quantified the frequency of positive BrdU calls by computing the z-score of positive calls against a binomial distribution as before. This allowed windows to be assigned as having either high or low BrdU signal (**Fig. 3e**). For example, two early firing replication origins on chromosome VI both give rise to BrdU positive regions on a single >100 kb read (read 3 in **Fig. 3e**), indicating that both origins activated in a single cell.

To explore this more widely, we visualised the BrdU frequency z-scores for individual nascent-strand reads that spanned known replication origins (each read considered covered >4kb either side of the origin location)^19^. In **Fig. 3f**, each row represents an individual nanopore sequencing read centred upon an origin. The colour gradient indicates the frequency of BrdU calls within 2 kb windows, and reads are sorted vertically by the population average BrdU-seq data: reads that span the most active origins will be near the top. In the majority of reads, we observed the highest frequency of positive BrdU calls at the origin. Most of these nascent molecules span efficient, early activating origins (‘unchecked’ by the intra-S phase checkpoint) indicative of origin firing during the BrdU pulse; there are only occasional examples of molecules where we detect BrdU incorporation at less efficient, late activating origins (‘checked’ by the intra-S phase checkpoint). Computing the average for each column the BrdU signal for unchecked (or checked) origins showed that on average early activating origins had incorporated more BrdU than later activating origins; in both cases the signal was symmetric and centred on the origins (**Fig. 3g**). However, some individual molecules showed asymmetric levels of BrdU incorporation relative to the origin, indicative of different rates of sister fork progression (**Supplemental Fig. S8**). We note that the BrdU signal in individual molecules is highest at replication origins and falls away as a function of distance from the origin (**Fig. 3e** and **f**). Eventually, the frequency of positive BrdU calls in a window drops below our detection threshold (z-score = −2, see **Fig. 3b**) and is called as a thymidine window. This implied a time-dependent drop in BrdU incorporation as the fork progressed away from the origin. This led us to hypothesize that we might be able infer fork-direction from the gradient of BrdU signal.

### Single-molecule replication dynamics

To examine replication dynamics in the absence of replication stress, we synchronised cell in G1 and released in the presence of a limiting concentration of BrdU (**Fig. 4a** and **Supplemental Fig. S7**). Samples were collected for D-NAscent, mass spectrometry, BrdU-seq, and DNA copy number measurements^20^. Using the D-NAscent results, we again summed all positive BrdU calls across all reads in non-overlapping 100 bp windows and found good agreement (Pearson correlation coefficient, *R*=0.75) with population-level BrdU-seq data (**Fig. 4b**); both assessments confirmed that the population average level of BrdU incorporation was inversely correlated with average replication time (**Supplemental Fig. S9**). Within individual reads, we assessed non-overlapping 2 kb windows and computed the z-score of positive BrdU calls in each window to a binomial distribution (examples shown in **Fig. 4c**). We observed clear peaks of BrdU incorporation at locations near known replication origins. The extremely long nanopore sequence reads allowed the identification of multiple active origins on single molecules. The BrdU signal either side of each origin declines, indicative of the progression of bi-directional replication forks at a time when the cellular BrdU concentration is falling. As forks move further away from initiation sites, the frequency of positive BrdU calls drops below our detection threshold (red to blue in **Fig. 4c**). Therefore, we consider the level of BrdU signal as a measure for when a sequence replicated; given that replication fork velocity is ∼2 kb/min^4, 21^ and that we observe differences in BrdU incorporation at 2 kb resolution, this indicates that we have close to 1 min temporal resolution.

**Figure 4:**
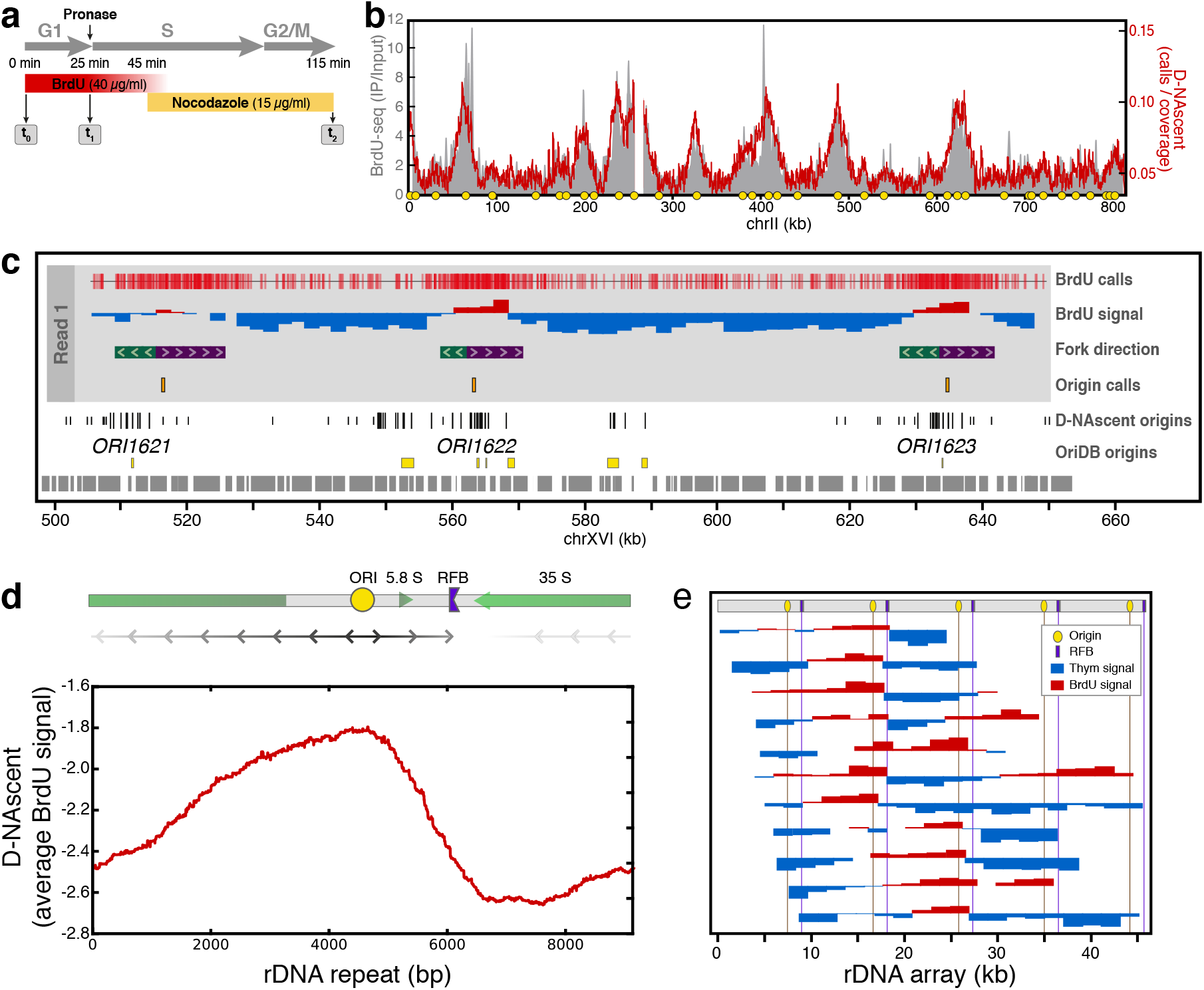
Single-molecule detection of replication dynamics. (**a**) Schematic of the experimental strategy for detection of replication dynamics by D-NAscent. At the indicated timepoints, samples were taken for mass spectrometry, DNA copy number measurements, BrdU-seq and D-NAscent. (**b**) Comparison BrdU-seq data and an ensemble of D-NAscent data across chromosome II (from timepoint t2). (**c**) An example 160 kb nanopore sequencing read showing BrdU calls at individual 6-mers (top track), the z-score for each 2kb window where BrdU-positive window scores are shown in red and thymidine-only window scores are shown in blue (middle track), and called fork direction and replication initiation sites (lower tracks). Origin calls from all spanning nanopore reads (black bars: tall, close to known origins; short, >3.9 kb (**Supplemental Fig. S10**) from known origins) and origins annotated as confirmed or likely by OriDB (yellow boxes) are displayed. (**d**) (*top*) A schematic representation of a single rDNA repeat showing the origin, replication fork barrier (RFB), predominant replication fork direction (line arrows) and the major transcripts (open arrows). (*bottom*) An ensemble of D-NAscent z-scores averaged over all nanopore sequence reads that spaned an rDNA repeat and had at least one BrdU-positive 2kb window. (**e**) The D-NAscent BrdU signal from selected molecules aligned to multiple rDNA repeats (origin, yellow; RFB, purple).

We determined the gradient of BrdU signal as a proxy for replication fork direction across all nascent strand sequence reads (examples shown in **Fig. 4c**). We used regions of diverging replication forks to call sites where replication initiated early in S phase prior to the BrdU concentration dropping below our detection threshold. This provides a whole genome map of DNA replication origin activity on single molecules. Examining all identified replication initiation sites revealed two distinct classes (**Supplemental Fig. S10**). The first class were found on multiple independent sequence reads, consistent with high efficiency origins used in many cells. These sites corresponded to replication origins identified in population level analyses^19^. The second class were dispersed throughout the genome with each site identified in a minority of molecules. These sites did not correspond to known replication origins. We considered that these sites might not represent genuine bi-directional replication initiation, for example, they could represent BrdU incorporation from a DNA repair pathway prior to S phase or a sequencing/analysis artefact. However, several lines of evidence argue in favour of these sites representing genuine replication origins. First, DNA repair synthesis prior to S phase would be confined to parental strands and synthesis would be unidirectional; the novel initiation sites we identified are present on nascent strands and show bi-directional synthesis (example shown in **Fig. 4c**). Second, applying more conservative criteria for origin identification (a higher z-score and a longer contiguous region of BrdU-positive 2kb windows) reduced the number of origin calls, but did not diminish the proportion of initiation events at novel locations (data not shown). Third, the pattern of BrdU detected at novel initiation sites closely resembled that observed at previously reported origins. Therefore, the single-molecule resolution of D-NAscent allows the detection of replication initiation sites that are too infrequently used to be detected by population-level methods.

### Identification of replication fork pausing on single-molecules

Detection of BrdU incorporation differences across nanopore sequencing reads allowed us to infer the relative replication time at ∼2 kb or ∼1 min resolution. Next, we sought to determine whether this could allow the detection of replication fork pausing events. The yeast ribosomal DNA (rDNA) repeats each contain a replication origin and a programmed unidirectional replication fork barrier (RFB) that pauses one of the sister forks (**Fig. 4d**)^22^. The repetitive nature of the rDNA limits their analysis by short-read technologies, but we were able to sequence thousands of molecules that each spanned multiple repeats. An ensemble of D-NAscent data analysed across a single rDNA repeat clearly demonstrated an asymmetric peak in the BrdU average z-score signal (**Fig. 4d**). In this ensemble analysis, the population average BrdU signal is maximal at the replication origin. The dramatic fall in signal to the right of the origin indicates a substantial delay to the progress of the rightward moving fork. This delay is positioned over the RFB and is consistent with unidirectional fork pausing. By contrast, the leftward moving fork shows no such delay. Analysis of single molecules (**Fig. 4e**) demonstrates firing of the origin in a subset of repeats and pausing of rightward moving forks at the RFB. Therefore, time-dependent reductions in BrdU incorporation frequency allow comprehensive analysis of replication dynamics on single molecules, revealing fork direction, initiation sites, termination sites and fork pausing/stalling.

## Discussion

We have developed a genomic single-molecule method for the detection of base analogues that we term D-NAscent. Base analogues are widely used in modern biology for the study of chromosome biology^23^, cell proliferation^24^ and gene expression^25^. Therefore, D-NAscent offers a powerful method for the advancement of each of these fields. Key features of nanopore sequencing^26^ make D-NAscent possible: the lack of an obligatory amplification step ensures that *in vivo* incorporated analogues are present in the sequenced strand; the interrogation of single nucleic acid strands permits direct detection of the analogue and provides single molecule information; and the extremely long sequence read lengths (>100 kb) allow detection of long-range *cis* interactions. We demonstrate that the currently available ONT MinION nanopore sequencing platform gives robust detection of thymidine analogues across the full range of sequence contexts (**Fig. 1** and **Fig. 2**). This allowed us to develop a model for the detection of *in vivo* incorporated BrdU that we validate against mass spectrometry and population level BrdU-seq data. The sensitivity of our BrdU-detection model allows us to measure changes in BrdU incorporation frequency on nascent strands and thereby reveal the temporal order of DNA replication on single molecules.

DNA replication is a stochastic process and many aspects, including replication origin activity, are masked in population-based approaches. Historically, this has required the use of complex, low-throughput, and low-resolution methodologies to visualize DNA replication on single-molecules. By applying D-NAscent to the study of yeast chromosome biology, we have generated the first whole genome map of DNA replication dynamics at the single molecule level. Unexpectedly, we discovered a novel class of replication origin that could not have been discovered by established methods. While most initiation sites that we detected (∼80%) are near known origins, approximately one fifth of replication initiation events occur at sites dispersed throughout the genome. Yeast replication origins were first characterized by their ability to support plasmid replication (as autonomously replicating sequences, called ARS elements)^27^ and it was subsequently shown that the same sequences can support replication initiation at their endogenous chromosomal locations^28^. Neither the plasmid nor the chromosomal assays have the sensitivity to detect very low efficiency origins. Recent *in vitro* studies have demonstrated origin-dependent and independent DNA replication initiation, due to promiscuity in the binding of the origin recognition complex^29, 30^. Although the *in vitro* origin-independent replication initiation was only observed in the absence of physiological levels of competitor DNA, it is consistent with our finding that many genomic locations can function at low frequency as a replication origin. Thus, we propose that replication of the yeast genome is initiated from both well-defined high-efficiency origins and a broadly distributed set of very low-efficiency origins, similar to the configuration observed in mammalian cells^2^.

The D-NAscent single-molecule methodology will allow many unresolved questions in chromosome biology to be addressed. For example, the single-molecule nature will allow the identification of *cis* regulatory mechanisms. The power to explore *cis* regulatory mechanisms is enhanced by the extremely long sequencing reads; in this study we present reads <160 kb, but others have reported ultra-long reads of >1 Mb. As such, D-NAscent complements recently developed single cell approaches for the study of DNA replication^31, 32^. Single cell approaches have relatively low spatial resolution, but they can provide *trans* information missing in single molecules. However, we and others have discovered that replication origin activity is generally regulated in *cis*^33-36^ emphasising the importance of the single molecule approach. Second, variants of the MinION sequencing method allow capture of sequence information from both DNA strands (1D^2^). Combining D-NAscent with 1D^2^ sequencing has the potential to reveal sites of conservative DNA replication, for example during recombination-dependent DNA synthesis^37, 38^. Third, extremely long sequencing reads allows D-NAscent to examine patterns of DNA replication in complex genomic locations (e.g. non-unique or repetitive sequences; **Fig. 4d** and **4e**) that are abundant in mammalian genomes and generally understudied. Fourth, the ability of D-NAscent to detect nascent DNA depends on the incorporation of thymidine analogues; achievable in many organisms and all commonly utilised model systems. This, and the gigabase throughput of nanopore sequencing platforms will allow the application of D-NAscent to many organisms, including the study of large stochastically replicated mammalian genomes. Existing single-molecule methods, such as DNA combing, have revealed extensive variability in replication initiation site usage and in fork progression rates. However, combed molecules are generally anonymous precluding the identification of chromatin features associated with variable fork velocity or replication initiation. Therefore, D-NAscent will allow the genome-wide identification of mammalian replication origins.

## Acknowledgments

We are grateful to John Diffley and Philippe Pasero for kindly providing strains; Amanda Williams and Rebecca Busby for Illumina NextSeq support; Michal Maj for assistance with flow cytometry; Joseph Caesar for IT infrastructure support; Microsoft and NVIDIA for computing resources. We thank Nieduszynski group members for helpful discussion and advice; Anthony Carr, Anne Donaldson, Shin-ichiro Hirage and David Sherratt for critical reading of the manuscript.

This work was supported by Biotechnology and Biological Sciences Research Council grant BB/N016858/1 and Wellcome Trust Investigator Award 110064/Z/15/Z. CAM is a Queen’s College Extraordinary Junior Research Fellow in Physiology; MAB is a St. Cross College Emanoel Lee Junior Research Fellow.

## Author contributions

CAN, CAM and MAB designed the study; CAM designed and performed the experiments; MAB designed and implemented the analogue-training and detection software; CAM and MAB undertook data analysis; PS undertook mass spectrometry, supervised by BMK and SK; CAN and JTS supervised the study; CAN, CAM and MAB wrote the paper.

## Conflict

JTS receives research funding from Oxford Nanopore Technologies and has received travel support to speak at meetings organized by Oxford Nanopore Technologies.

## Methods

### Defined substrates

Primers CA1218 and CA1219 (**Supplemental Table S2**) were annealed and extended with BIOTAQ DNA polymerase (Bioline) in the presence of dCTP, dGTP, dATP and either dTTP, BrdU-TP, FldU-TP, IdU-TP or EdU-TP (Jena Bioscience) each at 5 mM. Nanopore substrates must exceed a length of 250 bp. Thus, extended primers were digested with *Xma*I (NEB) and ligated to *Age*I-digested DNA sequences (>350 bp). Ligation products were gel purified prior to Nanopore sequencing.

### Yeast DNA for model training

Thymidine-auxotrophic yeast strain YLV11 was grown overnight in YPG (Formedium) supplemented with 100 μM Thymidine. Cells were then diluted to an OD_600_ of 0.06 into fresh YPG supplemented with 100 μM of BrdU and/or Thymidine (0%, 40%, 60%, 80% or 100% BrdU). Cells were grown at 30°C for 24 hours before samples were taken for Nanopore sequencing, MS analysis and BrdU-IP sequencing.

### Yeast cell cycle experiments

Cell cycle experiments were performed with yeast strain E3087 (**Supplemental Table S3**)^39^. Cells were grown in YPD media and arrested in G1 phase using alpha-factor. BrdU was added to a final concentration of either 400 μg/ml (hemi-labelled genomic DNA relating to Figure 3a) or 40 μg/ml (HU experiment and limiting BrdU concentration experiment, Figures 3c-f and 4, respectively). BrdU was added 25 minutes prior to Pronase-mediated release into S phase, followed by an arrest in G2/M by nocodazole treatment. For the HU experiment, 200 mM HU were added concomitantly with BrdU; cells were released into S phase for 45 minutes, then 400 μg/ml Thymidine was added and 15 minutes later, cells transferred into fresh YPD with Thymidine (400 μg/ml). Flow cytometry samples were taken at regular time intervals to assess cell cycle progression of each time course. Samples were treated with RNAseA and ProteinaseK prior to DNA staining with SYTOX Green (ThermoFisher S7020) and analysis on a BD LSRFortessa X-20 cell analyser. Samples for Nanopore sequencing, MS analysis, DNA copy number measurements and BrdU-IP sequencing were taken at defined time points in every cell cycle experiment. Genomic DNA was purified using phenol-chloroform extraction, RNAseA and PK treatment followed by ethanol precipitation.

### Mass spec validation

The equivalent ratio of 1 μg of DNA in 100 μl of water was added to 200 μl of hydrolysis solution (100 mM NaCl, 20 mM MgCl2, 20 mM Tris pH 7.9, 1000 U/ml Benzonase, 600 mU/ml Phosphodiesterase I, 80 U/ml Alkaline phosphatase, 36 μg/ml EHNA hydrochloride, 2.7 mM deferoxamine). The mixture was incubated for two hours and then lyophylised by SpeedVac. The lyophylisate was resuspended in 100 μl of buffer A per 1 ug of DNA used and half was transferred into an LC-MS vial for analysis. For the analysis by HPLC– QQQ mass spectrometry, a 1290 Infinity UHPLC was fitted with a Zorbax Eclipse plus C18 column, (1.8 μm, 2.1 mm 150 mm; Agilent) and coupled to a 6495a Triple Quadrupole mass spectrometer (Agilent Technologies) equipped with a Jetstream ESI-AJS source. The data were acquired in dMRM mode using positive electrospray ionisation (ESI1). Mass spectrometry was used for rare nucleosides and abundant nucleosides were quantified by HPLC-UV. The AJS ESI settings were as follows: drying gas temperature 230 °C, the drying gas flow 14 lmin-1, nebulizer 20 psi, sheath gas temperature 400 °C, sheath gas flow 11 lmin-1,Vcap 2,000 V and nozzle voltage 0 V. The iFunnel parameters were as follows: high pressure RF 110 V, low pressure RF 80 V. The fragmentor of the QQQ mass spectrometer was set to 380 V and the delta EMV set to +200. The UV quantification wavelength was 254 nm. The gradient used to elute the nucleosides started by a 5-min isocratic gradient composed with 100% buffer A (10 mM ammonium acetate, pH 6) and 0% buffer B (composed of 100% methanol) with a flow rate of 0.400 ml min-1 and was followed by the subsequent steps: 5-8 min, 94.4% A; 8–9 min, 94.4% A; 9– 16min 86.3% A; 16–17 min 0% A; 17– 21 min 0% A; 21–24.3 min 100% A; 24.3–25min 100%A. The gradient was followed by a 5min post time to re-equilibrate the column. The raw mass spectrometry data was analysed using the MassHunter Quant Software package (Agilent Technologies, version B.07.01). For the identification of compounds, raw mass spectrometry data was processed using the dMRM extraction function in the MassHunter software.

### Illumina population data

Yeast genomic DNA samples were assessed by BrdU-seq using the NextSeq 500 (Illumina). Genomic DNA was sheared to ∼300 bp using a Bioruptor. Sheared DNA was end-repaired and A-tailed using the NEBNext Ultra II end-repair module (E7546). A-tailed genomic DNA was barcoded using Illumina-compatible primers and NEBNext Ultra II ligation mix. Equal quantities of barcoded DNA samples were pooled and 20 ng of pooled DNA was reserved as “Input”. At least 1 microgram of pooled barcoded DNA was denatured and subjected to immuno-precipitated using an anti-BrdU antibody (BD 347580) and Protein-G dynabeads (ThermoFisher 10003D). Immuno-precipitated DNA was purified with AMPure XP bead. The immuno-precipitated and input DNA samples were PCR amplified separately using Illumina-compatible indexing primers and the NEBNext Ultra II Q5 Master Mix. DNA samples were sequenced (80 bp single-end) on a NextSeq 500. Illumina sequencing reads were demultiplexed and the barcode sequences were trimmed using the FASTX toolkit. Sequencing reads were mapped to the sacCer3 genome assembly using bowtie2. Read tag counts were determined for the 5’ end of uniquely mapping reads without mismatches in 100 bp nonoverlapping regions. The ratio between IP and Input sample was calculated for each bin, excluding bins that had less than 20% of the expected number of reads in the input sample. Ratios were median-smoothed over 1 kb windows.

### Nanopore sequencing

Samples were prepared for nanopore sequencing according to recommendations by Oxford Nanopore Technologies (ONT). The 2D library kit (SQK-LSK208) and 1D^2 library kit (SQK-LSK308) were used for synthetic substrates (Fig 1), the 1D Ligation-based library kit (SQK-LSK109) was used for the HU (Fig 3c-f) and BrdU-depletion experiments (Fig 4), and the 1D Native barcoding genomic kit (EXP-NBD103 and SQK-LSK108) for yeast genomic training material (Fig 2 and Fig 3a,b). The yeast genomic training DNA was sheared to an average length of 8 kb using g-TUBE (Covaris, 520079). The input DNA for all other nanopore libraries was unsheared. Quantitities of input DNA were adjusted to average molecule lengths, ranging between 12 ng and 5 μg for short synthetic and unsheared high-molecular weight genomic DNA, respectively. Input DNA for all libraries was end-repaired using NEBNext End Repair Module (NEB, E6050). In addition, genomic input DNA was treated with NEBNext FFPE RepairMix (NEB, M6630) to repair nicks. End-repaired samples were purified using 1x (synthetic and genomic training material) or 0.4× (HU and BrdU-depletion experiment) AMPure XP beads (Beckman Coulter, A63880). Next, ONT barcodes and/or adaptors specific to each library kit are ligated onto the samples. For 1D libraries, adapters were ligated using NEBNext Quick T4 DNA Ligase (NEB, E6065), followed by library purification using 0.4× AMPure XP beads with ONT’s wash buffer enriching for long molecules. For pooled 1D libraries, end-repaired samples were first ligated to ONT barcodes using Blunt/TA Ligase Master Mix (NEB, M0367), cleaned up using 1x AMPure XP beads and pooled in equal amounts prior to adapter ligation and final purification as above. The 1D^2 library preparation included Adapter ligation using Blunt/TA Ligase Master Mix, 0.4× AMPure XP bead purification, followed by sequencing adapter ligation using Blunt/TA Ligase Master Mix and AMPure XP bead clean up with proprietary ONT ABB wash buffer. 2D library preparation included ligation using Blunt/TA Ligase Master Mix and a proprietary mix of two adapters, one linear, the other a biotinylated hairpin. For purification after adaptor and tether ligation, My-One Streptavidin C1 Dynabeads (Thermo Fisher) were used to enrich for molecules with a hairpin. Nanopore libraries were sequenced on a MinION Mk1 sequencer using flow cell versions R9 (2D library), R9.4 (1D ligation libraries) and R9.5 (1D^2 library).

### Model training

Nanopore reads were basecalled using the Albacore basecalling software (v2.1.10) provided by Oxford Nanopore Technologies. We aligned the reads to the *S. cerevisiae* sacCer3 genome assembly using minimap2 (v2.10) with the “-a map-ont” setting ^40^. We eliminated those reads that aligned to mitochondrial DNA or ribosomal DNA, and for each remaining read that mapped uniquely to the genome (mapping quality >= 20), we aligned the signal events to their respective positions on the reference using nanopolish eventalign. For each thymidine-containing 6mer in our reads, we gathered all signal events that aligned to that 6mer; hence, the events gathered for each 6mer were taken from a range of genomic sequence contexts. For each of these 6mers that had greater than 200 aligned events, the signal events were filtered for outliers using a DBSCAN algorithm to eliminate trace alignment artefacts and the remaining events were used to fit a bimodal Gaussian mixture model. For each component of the fit mixture model, we computed the KL-divergence against the ONT 6-mer pore model. The distribution that had the higher KL-divergence against the ONT pore model was designated as the BrdU distribution.

### BrdU detection

As in model training, Nanopore reads were basecalled using the Albacore basecalling software (v2.1.10) and the reads were aligned to the *S. cerevisiae* sacCer3 genome assembly using minimap2 (v2.10) on the “-a map-ont” setting. We found that incorporation of BrdU into nanopore reads disrupts the accuracy of Albacore basecalling (data not shown) so we designated the true sequence of the read to be the subsequence of the reference that the read aligned to.

Signal events were aligned to positions on the Albacore basecall using adaptive banded dynamic programming^41^. This allowed us to use our trained BrdU pore model in the alignment to account for the presence of BrdU in the sequence while also circumventing the high space and time complexities that can result from dynamic programming-based alignment approaches. With an alignment of events to the Albacore basecall, we then aligned the events to positions on the reference using the minimap2 alignment. We used this alignment to find the signal events that corresponded to each position on the subsequence of the genome that the read mapped to. We only attempted to make a BrdU call at 6mers for which the KL-divergence between the BrdU distribution and the ONT thymidine-only distribution was greater than two. For each of these 6mers in the aligned reference subsequence, we computed the log-likelihood that this 6mer contained at least one BrdU by building a hidden Markov model (HMM) for the subsequence consisting of the 6-mer of interest and the surrounding 20 base pairs (**Supplementary Fig. S5**). Each match state for this surrounding sequence was given the distribution from the ONT pore model corresponding to the 6-mer at that position. We used the forward algorithm to calculate the probability of the events aligned to this 41-mer when the match state at the central position was set to the ONT model distribution (thymidine only) and again when the match state at the central position was set to our trained BrdU distribution. Taking the log-ratio of these two probabilities specifies the log-likelihood of BrdU at this position. We considered positions where the log likelihood of BrdU exceeded 2.5 to be positive BrdU calls.

### Region calling and fork direction

Using the detection output, for each non-overlapping 2kb window, we computed both the number of positive BrdU calls (*k*) and the total number of sites where a call (BrdU or thymidine) was made (*n*). From the ROC curve analysis (see **Fig. 2f**) and the mass spectrometry results, we can compute the probability of making a positive BrdU call for one of the 6-mers in our trained BrdU pore model:

> *p* = true positive probability × fraction of thymidine substituted for BrdU.

A binomial distribution with parameters *n* and *p* gives a model for the expected frequency of positive BrdU calls if the window actually is BrdU-containing. We computed the z-score of making *k* positive calls from this binomial distribution: positive z-scores indicate that there are a high frequency of positive BrdU calls in the window while negative z-scores indicate that the frequency of positive BrdU calls is lower than expected for an average BrdU-positive window. We considered a window to be a region of BrdU incorporation if the z-score was greater than −2 (see **Fig. 3b**). Fork direction was determined by smoothing the z-scores across a read with a 10kb moving average filter and computing the central derivative of the z-score for each 2kb window. Windows that had a z-score greater than −2 (called as a BrdU window) and had a negative z-score derivative were classified as rightward moving fork windows, and windows that were called as BrdU and had a positive derivative were classified as leftward moving fork windows. Positions that had at a leftward moving fork region of at least 4kb to the left and a rightward moving fork region of at least 4kb to the right were called as replication initiation sites.

We determined the number of reads with BrdU positive windows from a substrate prepared from cells grown in the absence of BrdU. Of the 1100 reads assessed only five had a BrdU positive window. In each of these five reads only a single window was called as BrdU positive.

### Code availability

The D-NAscent software is available at https://github.com/MBoemo/DNAscent.git.

### Data availability

Raw and processed Illumina and MinION data are available from NCBI GEO under accession number xxxxxx.

